# Serotonergic plasticity in the dorsal raphe nucleus characterizes susceptibility and resilience to anhedonia

**DOI:** 10.1101/719286

**Authors:** Nandkishore Prakash, Christiana J. Stark, Maria N. Keisler, Lily Luo, Andre Der-Avakian, Davide Dulcis

**Affiliations:** Department of Psychiatry, University of California San Diego, La Jolla, CA 92093, USA

## Abstract

Chronic stress induces anhedonia in susceptible, but not resilient individuals, a phenomenon observed in humans as well as animal models, but the molecular mechanisms underlying susceptibility and resilience are not well understood. We hypothesized that the serotonergic system, which is implicated in stress, reward and antidepressant therapy, may play a role. We found that plasticity of the serotonergic system contributes to the differential vulnerability to stress displayed by susceptible and resilient animals. Stress-induced anhedonia was assessed in adult male rats using social defeat and intracranial self-stimulation (ICSS), while changes in serotonergic phenotype were investigated using immunohistochemistry and *in situ* hybridization. Susceptible, but not resilient, rats displayed an increased number of neurons expressing the biosynthetic enzyme for serotonin, tryptophan-hydroxylase-2 (TPH2), in the ventral subnucleus of the dorsal raphe nucleus (DRv). Further, a decrease in the number of DRv glutamatergic neurons was observed in all stressed animals. This neurotransmitter plasticity is dependent on DR activity, as was revealed by chemogenetic manipulation of the central amygdala, a stress-sensitive nucleus that forms a major input to the DR. Activation of amygdalar corticotropin releasing hormone (CRH)+ neurons abolished the increase in DRv TPH2+ neurons and ameliorated stress-induced anhedonia in susceptible animals. These findings show that activation of amygdalar projections induces resilience, and suppresses the gain of serotonergic phenotype in the DR that is characteristic of susceptible animals. This molecular signature of vulnerability to stress-induced anhedonia and the active nature of resilience could be a target of new treatments for stress-related disorders like depression.

**SIGNIFICANCE STATEMENT:** Depression and other mental disorders can be induced by chronic or traumatic stressors. However, some individuals are resilient and do not develop depression in response to chronic stress. A complete picture of the molecular differences between susceptible and resilient individuals is necessary to understand how plasticity of limbic circuits is associated with the pathophysiology of stress-related disorders. Using a rodent model, our study identifies a novel molecular marker of susceptibility to stress-induced anhedonia, a core symptom of depression, and a means to modulate it. These findings will guide further investigation into cellular and circuit mechanisms of resilience, and the development of new treatments for depression.

## INTRODUCTION

Anhedonia, or lack of interest or pleasure, is a debilitating symptom of several psychiatric and neurological disorders, including major depressive disorder (MDD) (American Psychiatric Association, 2013) that reflects impaired brain reward function. It has been modeled in rodents and primates using multiple behavioral paradigms that capture different aspects of reward processing (Treadway and Zald, 2011; Der-Avakian et al., 2015; Alexander et al., 2019).

The intracranial self-stimulation (ICSS) procedure provides a direct and quantitative measure of anhedonia in rodents (Olds and Milner, 1954; Carlezon and Chartoff, 2007; Negus and Miller, 2014), which can be induced by stressors like social defeat (Martinez et al., 2002b; Rygula et al., 2005; Der-Avakian et al., 2014; Riga et al., 2015). However, not all stressed animals develop anhedonia (Krishnan et al., 2007; Der-Avakian et al., 2014), a phenomenon mimicking resilience to stressful or traumatic experiences in humans (Zisook et al., 1997; Bonanno et al., 2002). The mechanisms determining susceptibility or resilience to stress-induced anhedonia have important implications for understanding and developing treatments for disorders like MDD (Russo et al., 2012).

Serotonergic transmission in the brain is implicated in the processing of stress (Chaouloff et al., 1999; Hale et al., 2012; Backström and Winberg, 2017), reward (Kranz et al., 2010; Seymour et al., 2012; Liu et al., 2014; Li et al., 2016; Wang et al., 2019) and emotional behaviors (Neufeld et al., 2002; Herrmann et al., 2006; Roiser et al., 2007). Its role in the pathophysiology of many neurological (Chugani et al., 1997; Ener et al., 2003) and psychiatric disorders (López-Ibor, 1988; Lucki, 1998; Muller and Homberg, 2015) is well established. Knockout mice lacking key components of serotonergic machinery show altered stress-related behaviors (Holmes et al., 2003; Lira et al., 2003; Adamec et al., 2006; Gutknecht et al., 2015). Extracellular levels of serotonin (Kawahara et al., 1993; Grønli et al., 2007; Mokler et al., 2007), expression of serotonin-related molecules (Adell et al., 1988; Zhang et al., 2012; Issler et al., 2014; Donner et al., 2018), serotonergic activity (Grahn et al., 1999; Paul et al., 2011; Grandjean et al., 2019) and innervation (Natarajan et al., 2017) have been shown to change in response to both acute and chronic stress. Such regulation is linked to behavioral changes including anhedonia (Berton et al., 1997; Wood et al., 2013; Lopes et al., 2016; Natarajan et al., 2017) and is altered by treatment with antidepressants and anxiolytics (Benmansour et al., 1999; Abumaria et al., 2007). Accordingly, we investigated serotonergic circuitry for mechanisms that could explain susceptibility and resilience to stress-induced anhedonia.

Serotonin (5-hydroxytryptamine or 5-HT) is mainly synthesized in the raphe nuclei (Dahlström and Fuxe, 1964; Steinbusch, 1981; Jacobs and Azmitia, 1992; Hornung, 2003) by the enzyme tryptophan hydroxylase 2 (TPH2) (Walther et al., 2003; Zhang et al., 2004). Among the raphe nuclei, the dorsal raphe nucleus (DR) accounts for most serotonergic cell bodies both in rat (Jacobs and Azmitia, 1992) and human brains (Baker et al., 1991; Baker et al., 1990). Furthermore, serotonergic transmission occurs at multiple sites in the central nervous system (Steinbusch, 1981; Jacobs and Azmitia, 1992; Vasudeva et al., 2011).

Earlier studies have found that the number of serotonergic neurons (Underwood et al., 1999), levels of TPH2 mRNA (Bach-Mizrachi et al., 2008) and TPH2 protein (Boldrini et al., 2005) are elevated in the DR of deceased patients who committed suicide. These results led us to hypothesize that chronic stress-induced plasticity of serotonergic expression (Demarque and Spitzer, 2010) may occur in the DR and play a role in determining susceptibility or resilience to anhedonia. Neurotransmitter plasticity (Dulcis and Spitzer, 2012; Dulcis, 2016) has not been explored previously in the DR or other brain regions in animal models of anhedonia.

Here, we describe a mechanism of transmitter plasticity involving increased number of TPH2+ neurons in the DR in susceptible rats following chronic stress. We further demonstrate that manipulation of DR activity can rescue the transmitter phenotype and behavior.

## METHODS

### Animals

Adult male rats were used for all experiments. All rats were housed in a 12h reverse light-dark cycle with *ad libitum* access to food and water. Wistar rats (Charles River Laboratories, Kingston, NY) weighing 300-400g (~8 weeks old) were used for ICSS electrode implantation surgeries, behavior and immunohistochemistry, in experiments without chemogenetic manipulation. *Crh-Cre* transgenic male Wistar rats (rat line generously provided by Dr. Robert O. Messing, University of Texas at Austin) were bred in our vivarium and used for the chemogenetics experiment. Details of development of the *Crh-Cre* rats are described by Pomrenze et al. (2015). For social defeat, male Long-Evans rats (retired breeders; Charles River Laboratories) co-housed with females and litters were used as resident aggressors. All rats were pair-housed except during social defeat. *Crh-Cre* breeding pairs were at least 10 weeks old and either pair-housed or harem-housed (two females with one male). Pups were weaned from the dam and genotyped 21 days after birth. All experiments were carried out in accordance with the guidelines of AAALAC International and National Research Council’s Guide for the Care and Use of Laboratory Animals and approved by the UCSD Institutional Animal Care and Use Committee.

### Genotyping

Ear tissue punches (2 mm diameter) were collected from *Crh-Cre* progeny for DNA extraction and genotyping. For DNA extraction, tissue was incubated in 75 μl alkaline lysis buffer (25 mM NaOH, 0.2 mM EDTA, pH 12.0) at 95°C for 1h followed by addition of equal volume of neutralization buffer (40 mM Tris-HCl, pH 5.0) and short-term storage at 4°C. The mixture was used as source DNA for PCR-based genotyping. PCR protocol: 0.5 μl of DNA, 0.5 μl each of forward and reverse primers, 5 μl of KAPA2G Fast HotStart ReadyMix (KK5603, KAPA Biosystems) and 3.5 μl of sterile water were mixed in a 10 µl reaction. Cre recombinase forward primer, 5’-GCATTACCGGTCGATGCAACGAGTGATGAG-3’ and reverse primer, 5’-GAGTGAACGAACCTGGTCGAAATCAGTGCG-3’ (Washington University Mouse Genetic Core, mgc.wustl.edu) were used. Cycling parameters were 95°C for 3’; 30 cycles of 95°C for 15’’, 60°C for 60’’, 72°C for 40’’; 72°C for 2’ in a T100™ Thermal Cycler (Bio-Rad) followed by long-term storage at −20°C. PCR product was analyzed by horizontal agarose gel electrophoresis and presence of 550 bp band was determined in order to identify Cre-positive progeny.

### Surgery

For ICSS electrode implantations, rats were anesthetized with a 5% isoflurane/oxygen vapor mixture and attached to a stereotaxic frame (Kopf Instruments; Tujunga, CA) where continuous flow of 2% isoflurane/oxygen was administered throughout the procedure. The incisor bar was set at 5.0 mm above the interaural line. Bipolar insulated stainless-steel electrodes (11 mm length, model MS303/2; Plastics One; Roanoke, VA) were unilaterally (counterbalanced) implanted in the posterior lateral hypothalamus (AP −0.5 mm, ML ± 1.7 mm from bregma and DV −8.3 mm from dura). The electrode was secured using dental acrylic and 4-6 stainless steel jeweler’s screws. The exposed electrode pedestal was shielded using a metal screw cap to prevent damage.

For chemogenetics, bilateral viral injections (1.0 μl/side) into the lateral subnucleus of the central amygdala (AP −2.3 mm, ML ± 4.7 mm from bregma and DV −6.9 mm from dura, head parallel to horizontal) were performed using a 30G metal cannula (PlasticsOne) connected to a Hamilton syringe pump (10 μl syringe) at a rate of 0.1 μl/min prior to electrode implantation during the same surgical procedure. Rats were injected with either the control virus (AAVDJ-Syn1-DIO-eGFP, 1.78E+13 GC/mL, Salk Institute; La Jolla, CA) or the excitatory DREADD receptor-encoding virus (AAV5-hSyn-DIO-hM3Dq-mCherry, 6.50E+12 GC/mL, Addgene, Watertown, MA). Viral incubation occurred for at least 8 weeks during the post-surgical recovery, ICSS training, baseline testing and saline habituation periods. Post-surgical treatment with topical antibiotic cream and 20 mg/kg of enrofloxacin IM was provided to prevent infection.

### ICSS apparatus, training, testing and analysis

The ICSS procedure, including apparatus, training and testing, was performed as previously described (Der-Avakian and Markou, 2010). Briefly, rats were trained to seek reinforcement of direct current stimulation of the posterior lateral hypothalamus by turning a wheel manipulandum in the testing chamber. A non-contingent stimulus (100 Hz electrical pulse train) of current intensity varying from 50 to 300 μA was delivered to the rat by means of a computer-controlled constant current stimulator (Stimtek Model 1200c; San Diego Instruments, San Diego, CA). Rats were trained to turn a wheel in response to the non-contingent stimulus to receive a second (contingent) stimulus with identical parameters. Current intensities were systematically varied across trials, separated by inter-trial intervals without stimulation. Using this discrete-trial current-intensity procedure, the minimum current intensity required to elicit a response from the rat was measured and defined as the reward threshold. Reward thresholds were measured every day, at the same time, in trained rats. Thresholds were monitored for 3 to 6 weeks until they were stable (i.e., <10% variation over 5 consecutive days). The baseline threshold for each rat was calculated as the average of its daily thresholds for 3 days prior to the beginning of testing. Rats with stable thresholds were divided into control and stress groups. For the chemogenetics experiment, litter effects were avoided by distributing rats from different litters across groups. Controls were tested in the ICSS procedure daily and stressed rats were tested daily within 15 minutes of a social defeat encounter. Elevations in thresholds indicated that a greater current intensity was required to generate positive reinforcement, reflecting an anhedonic state. A rat was classified as susceptible to anhedonia if its average threshold during days 19-21 of social defeat was greater than 3 standard deviations from the pre-defeat baseline thresholds of the cohort (calculated by averaging baseline thresholds across all stressed rats). Daily reward thresholds were plotted as percent changes from baseline averaged across rats in each group.

### Chronic social defeat

A resident-intruder procedure was used as previously described (Der-Avakian et al., 2014). Long Evans males (residents), pre-screened for aggression and dominance, were housed in a large cage (61 cm × 43 cm × 20 cm) with females and progeny. During social defeat (21 days), the experimental male Wistar (intruder) was co-housed with the resident but physically separated by an acrylic partition that allowed for exchange of visual, auditory and olfactory information between the intruder and the residents. For 3 min each day, the female resident and pups were removed from the cage, and the partition was lifted to allow a direct physical interaction between the males. Social defeat was defined as a supine submissive posture of the intruder for 3 consecutive seconds with the resident pinning the intruder down. After a social defeat encounter or 3 min (whichever occurred first), the intruder was removed and its reward threshold was measured. At the end of each day of testing, the intruder was paired with a different resident for the next 24 h period.

### Drug treatment

Clozapine (0.1 mg/kg; MP Biomedicals, Santa Ana, CA) was dissolved in 0.1% dimethyl sulfoxide (DMSO) in sterile saline and administered intraperitoneally (IP) once daily, 30 min prior to ICSS testing during the 21-day social defeat procedure. For habituation to IP injections, rats across all experimental groups were administered 0.1 ml of 0.1% DMSO in saline (equal volume and route of administration as for subsequent clozapine injections) 30 min prior to ICSS testing daily, for 7-14 days prior to the start of social defeat until their ICSS baseline thresholds stabilized.

### Tissue collection and processing

In all experiments, on day 21, 6 h after social defeat/ICSS testing, rats were administered 0.2-0.3 g/kg Fatal-Plus C IIN (pentobarbital sodium; IP). After complete loss of reflexes, rats were transcardially perfused with phosphate-buffered saline (PBS, pH 7.5) until perfusate was colorless, followed by perfusion with equal volume of ice-cold 4% paraformaldehyde (PFA, pH 7.5) dissolved in PBS. Rats were then decapitated and whole brains were extracted and post-fixed in fresh 4% PFA for 24 h at 4°C followed by incubation in 30% sucrose at 4°C for 48 h or until brains were completely submerged. Thirty μm coronal sections of VTA, DR and 50 μm sections of the hypothalamus and amygdala were collected using a microtome (Leica) in a cryoprotectant solution (30% v/v glycerol, 30% v/v ethylene glycol in PBS, pH 7.4) for long-term storage at −20°C to −40°C. For each brain region, every fourth section was collected in the same well as a set of tissue for a given staining procedure.

### Immunohistochemistry and *in situ* hybridization

Antibody details and concentrations are provided in Table 1. All washes and incubations were performed with gentle shaking. Antibodies were diluted in blocking solution (5% normal horse serum, 0.3% Triton-X 100 in PBS). For immunofluorescence, sections were washed 3 times for 5’ (3 × 5’) in PBS, incubated in blocking solution for 30’ to 1h, incubated in primary antibody solution overnight at 4°C, washed 3 × 10’ in PBS, incubated in secondary antibody solution for 1h at room temperature, washed 3 × 10’ in PBS, mounted on a positively charged glass slide (Fisherbrand Superfrost Plus) in 0.2% gelatin in PBS, coverslipped with mounting medium (Fluoromount-G^®^, SouthernBiotech) and sealed with nail polish.

**Table 1:** Antibodies. Primary and secondary antibodies used in this study listed with details of host and target species, type (polyclonal, monoclonal or secondary), working dilution and product supplier.

| **Antibody** | **Dilution** | **Catalog # (Manufacturer)** |
| --- | --- | --- |
| Mouse $\alpha$ cFos (monoclonal) | 1:1000 | sc-166940 (Santacruz Biotech) |
| Guinea pig $\alpha$ NeuN (polyclonal) | 1:2000 | ABN90 (Millipore Sigma) |
| Sheep $\alpha$ TH (polyclonal) | 1:1000 | AB1542 (Millipore Sigma) |
| Rabbit $\alpha$ TPH2 (polyclonal) | 1:500 | PA1-778 (Thermo Fisher Scientific) |
| Guinea pig $\alpha$ VGlut3 (polyclonal) | 1:100 | AB5421-I (Millipore Sigma) |
| Sheep $\alpha$ bNOS (polyclonal) | 1:500 | AB1529 (Millipore Sigma) |
| Donkey $\alpha$ sheep Alexa Fluor 647 (secondary) | 1:500 | A21448 (Invitrogen) |
| Donkey $\alpha$ sheep Alexa Fluor 488 (secondary) | 1:500 | A11015 (Invitrogen) |
| Donkey $\alpha$ guinea pig Alexa Fluor 488 (secondary) | 1:500 | 706-545-148 (Jackson ImmunoResearch) |
| Donkey $\alpha$ rabbit Alexa Fluor 555 (secondary) | 1:500 | A31572 (Invitrogen), SAB4600177 (Sigma) |
| Horse $\alpha$ rabbit biotinylated (secondary) | 1:300 | BA-1100 Vector Laboratories, CA |
| Donkey $\alpha$ mouse Alexa Fluor 647 (secondary) | 1:500 | A31571 (Invitrogen) |

For colorimetric DAB-based immunohistochemistry, brain sections were processed in the following steps: 3 × 5’ PBS washes, incubation in 3% hydrogen peroxide in PBS for 15’, blocking solution for 30’, primary antibody overnight at 4°C, 3 × 10’ PBS washes, secondary antibody incubation at room temperature, 3 × 10’ PBS washes, incubation in fresh ABC solution (1:1 mixture of Reagents A and B, each diluted 1:100 in 2% NaCl/0.3% Triton-X 100/PBS) (VECTASTAIN^®^ HRP Kit, Vector Labs), 3 × 10’ PBS washes, incubation for 3-5’ in 3,3’ diaminobenzidine (DAB, Acros Organics) staining solution (0.025% w/v DAB, 0.01% v/v hydrogen peroxide in PBS), 3 × 10’ PBS washes, mounting in 0.2% gelatin in PBS, drying and coverslipping in Cytoseal™ 60 (Thermo Scientific) and sealing with nail polish.

RNAscope^®^ v 2.0 (ACD Bio) *in situ* hybridization in combination with immunofluorescence was performed as per manufacturer’s instructions for *Tph2* (Part ID 316411), *Vglut3* (Part ID 476711-C2) and *Pet1* (*Fev*) (Part ID 487771-C3) mRNA transcripts.

### Image acquisition and processing

For fluorescence staining, multi-channel confocal z-stacks of each tissue section were acquired with 2.5 μm distance between optical sections using a Leica SPE confocal microscope (10X or 20X dry objective). Objective resolution and acquisition settings (laser power, gain, pinhole aperture and signal averaging) were applied uniformly across sections within a given experiment. Maximum intensity z-projections were made using Fiji (Schindelin et al., 2012) image analysis software. Linear brightness and contrast adjustments were applied uniformly across all pixels for each image. Images in TIFF format were used for quantification. Mean filter (Fiji) or box blur (Adobe Photoshop CS4) with 2-pixel radius was applied to representative images in figures.

For DAB-stained sections, 20X brightfield images were acquired using an Olympus Virtual Slide Microscope (VS120) and stored in TIFF format for quantification.

### Quantification of cell number

For quantification of cell numbers for each marker, fluorescence or brightfield images (processed as described above) were organized according to their rostrocaudal position and specific sections were chosen by visual inspection for quantification and analysis using the Rat Brain Atlas (Paxinos and Watson, 2014) as a reference. For quantification of TPH2, cFos, NeuN, VGLUT3, nNOS immunofluorescence and *Tph2*, *Vglut3* and *Pet1* mRNA in the mid-DR, sections at rostrocaudal positions −7.76, −7.88 and −8.00 mm from bregma were selected and subdivisions were demarcated based on TPH2 staining pattern with reference to Abrams et al. (2004), Kelly et al. (2011) and Paxinos and Watson (2014). For quantification of TPH2 using DAB staining, sections at rostrocaudal positions −7.64, −7.76, −7.88, −8.00, −8.12, −8.24 and −8.36 mm from bregma were chosen. Only cells that were in focus, with clearly discernible cellular appearance (size, shape and cell boundaries) and with intense colorimetric or fluorescent stain filling at least 50% of the cell’s area (by visual estimation), were considered positive. For quantification of TH in the VTA, every fourth 30 µm section between rostrocaudal positions −5.60 mm and −6.20 mm from bregma was chosen. VTA was demarcated based on TH expression with reference to Paxinos and Watson (2014). Fluorescently stained cell bodies were counted unilaterally in the parabrachial pigmented and paranigral nuclei of the VTA and doubled prior to analysis and plotting. For quantification of TH in the periventricular nucleus of the hypothalamus, every fourth 30 µm section between rostrocaudal positions −1.40 mm and −2.00 mm from bregma was chosen. The periventricular nucleus was identified by TH expression with reference to Paxinos and Watson (2014) and cell bodies were counted unilaterally and doubled prior to analysis and plotting. In all animals with chemogenetic manipulation, validation of DREADD receptor expression was performed by visually examining sections of the central amygdala (lateral subnucleus, bilateral) throughout its rostrocaudal extent, for native mCherry or eGFP expression under a fluorescence microscope.

### Experimental design and statistical analyses

IBM SPSS Statistics 24 software was used for all statistical testing. Reward threshold comparisons for non-chemogenetic experiments were made using a mixed ANOVA (with Greenhouse-Geiser correction for sphericity violation as ε<0.75) after testing for normality (Shapiro-Wilk’s test) and homogeneity of variance (Levene’s test). *Day* was included as a within-subjects factor and *Stress* (control, susceptible and resilient) was the between-subjects factor. Significant main and interaction effects were followed with Bonferroni *post hoc* tests. For cell quantification comparisons in non-chemogenetics experiments, a 1-way ANOVA was used (control, susceptible and resilient groups) after ensuring normality and homoscedasticity of data (Shapiro-Wilk’s test and Levene’s test, respectively) and Tukey’s HSD *post hoc* tests were performed if applicable. Data that did not satisfy criteria for an ANOVA were analyzed using Kruskal Wallis H tests and *post hoc* comparisons with Bonferroni correction were performed.

For the chemogenetics experiment, rats either expressed the GFP or hM3Dq virus and were split into control and defeat groups after their ICSS baseline thresholds stabilized. All rats were administered 0.1% DMSO in saline (vehicle) during habituation and clozapine during the 21d social defeat period. GFP- and hM3Dq virus expressing non-stressed controls were pooled after ensuring that there was no statistical difference between the groups. Effects of vehicle were tested using a 1-way repeated measures ANOVA (*Day* as within-subjects factor) that included all rats. Reward thresholds were compared using a mixed ANOVA with *Day* (for analysis of effects of vehicle and clozapine on control groups) or *Period* (acute: average of days 1-3 or chronic: average of days 19-21, for analysis of effect of clozapine on stressed groups) as the within-subjects factor and *Group* (Control, GFP susceptible, GFP resilient, hM3Dq susceptible and hM3Dq resilient) as the between-subjects factor. Significant main and interaction effects were followed with Bonferroni *post hoc* tests. Cell count comparisons between the same five groups were conducted using an ANOVA with Tukey’s HSD for *post hoc* comparisons. The coefficient of correlation between TPH2+ counts and reward thresholds, was calculated using bivariate Pearson correlation test.

Alpha level was set to 0.05 for all analyses. Appropriate sample sizes for each experiment were determined with standard Cohen’s d power analysis with target effect size set to 0.8 and alpha level to 0.05. Outliers within any group, determined using the median absolute deviation method (Iglewicz and Hoaglin, 1993) were excluded from statistical analyses. Microsoft Excel and GraphPad Prism 8.0.2 were used for generating plots.

## RESULTS

### Susceptible rats show elevated ICSS thresholds after chronic social defeat

Monitoring ICSS thresholds over 21-days of social defeat revealed that a subset of rats was susceptible to stress-induced anhedonia, while others were resilient (Fig. 1A). Stressed rats whose thresholds at the end of the 21-day period, were greater than 3 standard deviations from baseline, were classified as ‘susceptible’ and others as ‘resilient’. Independently, a k-means cluster analysis of thresholds at the end of social defeat (averaged over days 19-21), split the cohort of stressed rats into identical groups, with susceptible rats forming a separate cluster from resilient rats and unstressed controls (Fig. 1B, right). The same cluster separation was not present at the beginning of social defeat (averaged over days 1-3), indicating that susceptible/resilient phenotypes were not predictable early during social defeat, in response to acute exposure, but rather developed over time due to chronic stress (Fig. 1B, left). There were also no differences in baseline thresholds across groups (Fig. 1C) as determined by a Kruskal-Wallis H test (χ^2^(2)=1.233, *p*=0.540), indicating that baselines thresholds do not predict ICSS responses to subsequent social defeat. A 2-way mixed ANOVA using Greenhouse-Geiser correction (ε=0.379) revealed a main effect of *Stress* (F(2,25)=15.998, *p*=3.40E-5; Fig. 1A) but no significant interaction between *Stress* and *Day* (*p*=0.106). However, based on previous evidence that susceptible/resilient phenotypes develop over time (Der-Avakian et al., 2014) which was supported by our observations (Fig. 1A,B), we hypothesized that the behavioral difference between susceptible and resilient rats arises at an intermediate timepoint during the 21-day social defeat paradigm. Accordingly, we performed *post hoc* pairwise comparisons with Bonferroni correction, which revealed that susceptible rats had significantly elevated thresholds (*p*<0.05) relative to controls on days 2-10 and relative to both resilient and control animals from day 11 onwards (Fig. 1A), while resilient rats did not differ significantly from controls. While controls gained weight over time, all stressed rats showed a slight decrease in weight over the 21-day stress period, indicating that the metabolic effects of stress were similar across susceptible and resilient rats (Fig. 1D). Latencies to supine submissive posture during social defeat, a quantitative measure of stress exposure, were also similar between susceptible and resilient groups (Fig. 1E), indicating that resilient animals were not subjected to any less stress than susceptible rats. Rats across groups did not show a difference relative to controls in latency to respond to ICSS stimulation, indicating that social defeat did not differentially affect motor activity between susceptible, resilient, and control rats (Fig. 1F). Number of injuries during social defeat was also similar across susceptible and resilient rats (Fig. 1G), indicating that elevated thresholds in susceptible rats were likely not a function of immune responses to injury.

**Figure 1:**
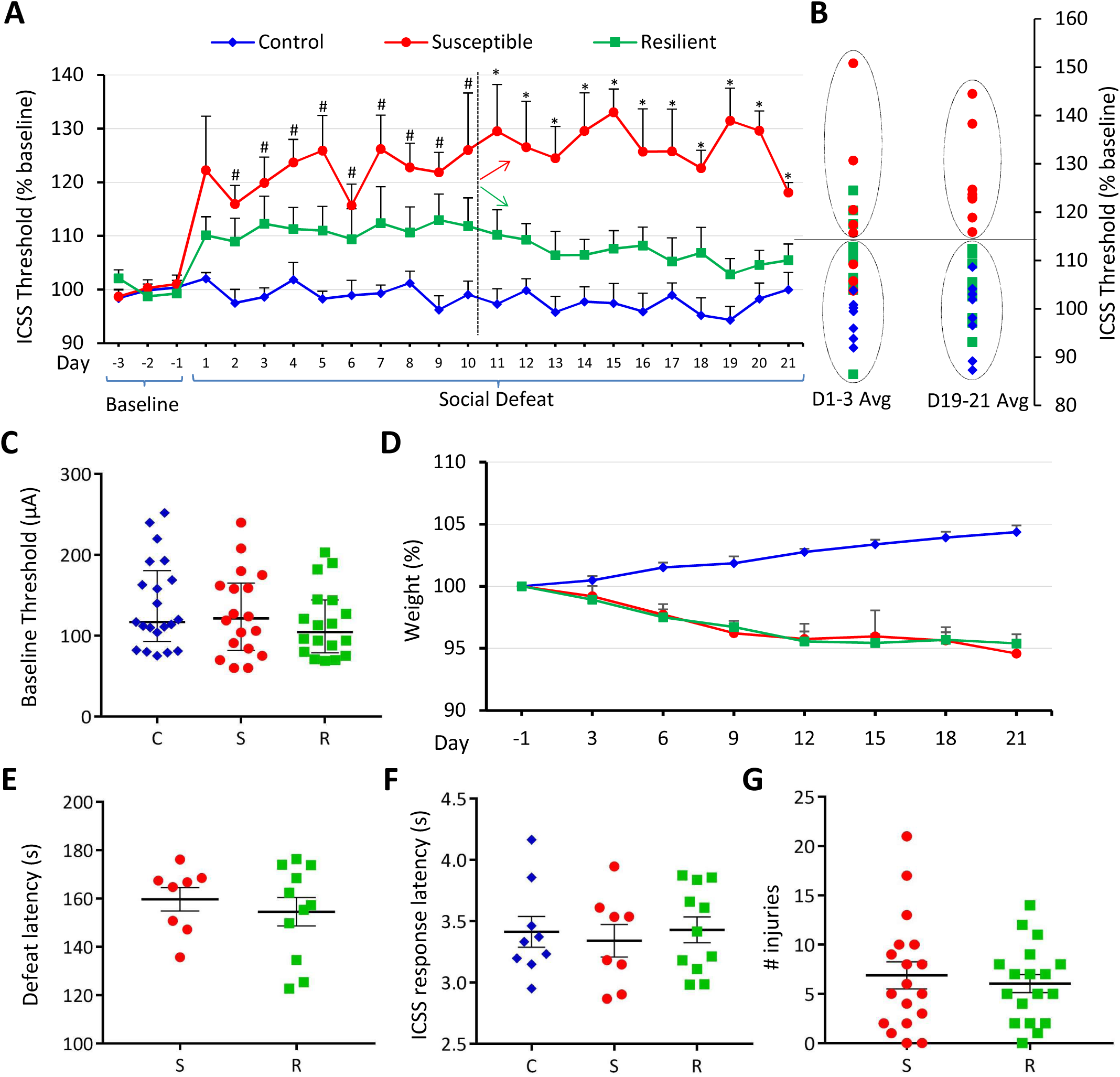
Susceptible rats show elevated ICSS thresholds during stress. **A**, Daily ICSS thresholds (mean across animals + s.e.m.) plotted as percent of baseline for 3 days prior to defeat and each day for 21-day social defeat. Control rats (*n=9*) represented by blue diamonds, susceptible rats (*n=8*) by red circles and resilient rats (*n=11*) by green squares. Significant (*p*<0.05) post-hoc pairwise comparisons for each day shown above error bars. # indicates significant difference between susceptible and control groups but not between other pairs. * indicates significant difference between susceptible and control as well as susceptible and resilient groups. Dashed vertical line after Day 10 indicates the day from which susceptible (red arrow) and resilient (green arrow) groups differ significantly. **B**, K-means cluster analysis of Days 1-3 (left) and Days 19-21 (right). **C**, Absolute baseline ICSS current intensity thresholds (in µA) for each experimental group (C, controls; S, susceptible; R, resilient). Values are median and interquartile range. **D**, Rat body weight (mean ± s.e.m.) in grams, measured every 3 days over 21-day period and plotted as percent change relative to baseline (Day -1) for each group. **E**, Latency to supine submissive posture during social defeat, in seconds, plotted for stressed groups as mean ± s.e.m. **F**, ICSS response latency (mean ± s.e.m.) in seconds, plotted for each group. **G**, Total number of injuries (mean ± s.e.m.) suffered by each rat during 21-day social defeat plotted for stressed group. For **B-E**, graph markers indicate experimental conditions as defined in **A**.

### Susceptible rats display an increased number of TPH2+ neurons in the DRv

Altered numbers of TPH2+ neurons in the DR have been observed in human victims of suicide (Underwood et al., 1999). Therefore, we counted the number of TPH2+ neurons in each of the following subnuclei of the mid-rostrocaudal DR: dorsal (DRd), ventral (DRv), ventrolateral “wings” (DRvl) and interfascicular (DRi) (Fig. 2A). The number of TPH2+ neurons in the DRv was significantly elevated (Kruskal Wallis H test, χ^2^(2)=11.528, *p*=0.003) in susceptible rats, relative to control (*p*=0.008, Bonferroni’s adjustment) and resilient rats (*p*=0.008, Bonferroni’s adjustment) (Fig. 2B,C). The DRd, DRvl and DRi showed no differences in the average number of TPH2+ neurons per section across groups as determined by Kruskal Wallis H tests (Fig. 2C). Independent Kruskal Wallis H tests with *post hoc* pairwise comparisons using Bonferroni adjustment revealed a significantly increased number of TPH2+ neurons (*p*<0.05) in susceptible rats relative to resilient and control rats at each of the seven rostrocaudal positions examined (Fig. 2D), indicating that stress-induced TPH2 expression was not localized to particular rostrocaudal sub-regions of the DRv. Since neurotransmitter plasticity involving dopaminergic neurons (marked by tyrosine hydroxylase, TH) has been observed in other brain regions, such as the periventricular nucleus (PeVN) of the hypothalamus after photoperiod stress (Dulcis et al., 2013) and the ventral tegmental area (VTA, Romoli et al., 2019) after neonatal exposure to nicotine, we investigated whether the number of TH+ neurons differed across our experimental groups in these regions. Neither of these regions showed differences across groups as determined by Kruskal Wallis H tests (Fig. 2E-H), indicating the specificity of TPH2 plasticity in the DR following social stress.

**Figure 2:**
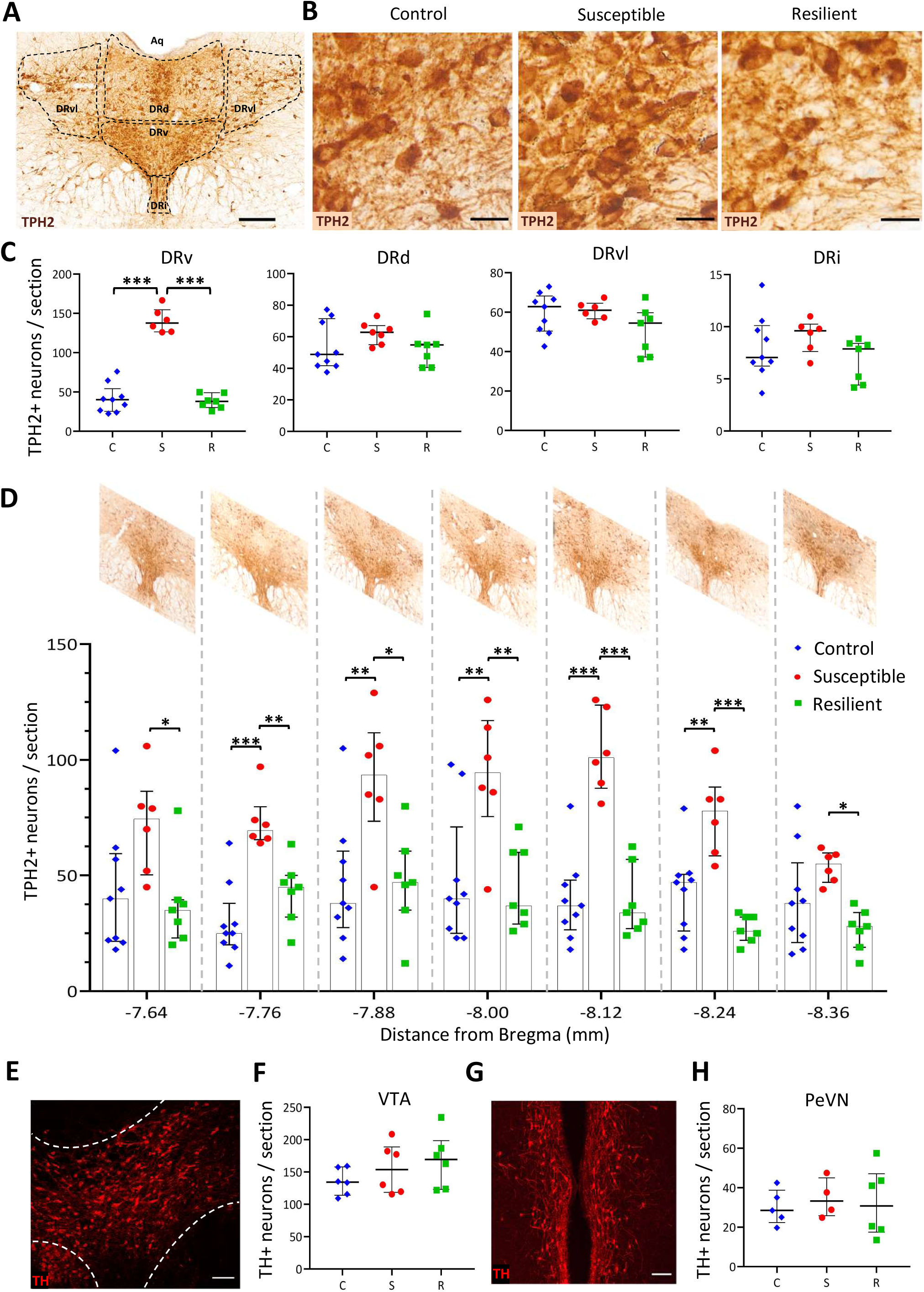
Susceptible rats display more TPH2+ neurons in the ventral subnucleus of the dorsal raphe nucleus (DRv) **A**, Representative coronal section through the dorsal raphe nucleus (DR) stained for tryptophan hydroxylase isoform 2 (TPH2, brown) by DAB immunohistochemistry. Various DR subnuclei present at the mid-rostrocaudal level are outlined in black. DRd, dorsal; DRv, ventral; DRvl, ventrolateral; DRi, interfascicular; Aq, aqueduct of Sylvius. Scale bar: 200 µm. **B**, Representative images of DRv sections stained for TPH2 from each experimental group. Scale bars: 25 µm. **C**, Number of TPH2+ neurons in ventral (DRv), dorsal (DRd), ventrolateral (DRvl) and interfascicular (DRi) subnuclei of the DR. Counts (per section) were averaged across rats (Controls, C, blue diamonds, *n=9*; susceptible, S, red circles, *n=6*; resilient, R, green squares, *n=7*) and plotted as median with interquartile range. *** *p* < 0.001. **D**, Quantification of number of TPH2+ neurons in the DRv at 120 µm rostro-caudal intervals in the middle DR. Rostro-caudal positions represented on x-axis as distance from bregma in mm and representative images of the DR at each position are shown above. Counts were averaged across rats (graph symbols and sample sizes same as in **C**) and plotted as median and interquartile range. * *p* < 0.05, ** *p* < 0.01, *** *p* < 0.001. **E**, Representative coronal section through the ventral tegmental area (VTA) seen unilaterally, outlined in white, stained for tyrosine hydroxylase (TH, red) to mark dopaminergic neurons. Scale bar: 100 µm. **F**, Bilateral quantification of the number of TH+ neurons per VTA section. Bilateral counts were averaged across 6 rats per group and plotted as median and interquartile range. Graph symbols as defined in **C**. **G**, Bilateral view of a representative coronal section through the periventricular nucleus of the hypothalamus (PeVN), stained for tyrosine hydroxylase (TH, red) to mark dopaminergic neurons. Scale bar: 100 µm. **H**, Bilateral quantification of number of TH+ neurons per PeVN section. Counts were averaged across rats (Controls *n=5*, susceptible *n=4*, resilient *n=6*) and plotted as median and interquartile range. Graph symbols as defined in *C*.

### TPH2/VGLUT3 switching occurs in susceptible rats following chronic stress

We first asked if the increase in TPH2+ neurons came from an increased number of neurons in the DRv. No significant differences were observed in the total number of mature neurons (marked by NeuN) in the DRv, as measured by a 1-way ANOVA (Fig. 3A,B). To identify the reserve pool (Dulcis and Spitzer, 2012) of differentiated DRv neurons that is recruited to acquire TPH2 in susceptible rats, we examined DRv for expression of other neurotransmitters and asked whether the extent of their co-expression with TPH2 changed across experimental groups. Of the various neurotransmitters expressed in the DR (Fu et al., 2010), we chose to examine those previously implicated in stress and reward. Mesolimbic dopamine is a well-known mediator of reward (Nestler and Carlezon, 2006) and we investigated whether DR dopaminergic (TH+) neurons could display plasticity of TPH2 phenotype. TH+ neurons in the DR did not display any overlap with TPH2 expression and were located very rostrally within the DR, outside the region where increased TPH2+ neurons were observed (Fig. 3C), consistent with previous observations (Fu et al., 2010). Another candidate neurotransmitter in the DR associated with reward and depression is nitric oxide (Gholami et al., 2003; Dhir and Kulkarni, 2007; Zhou et al., 2011). Nitrergic neurons, marked by neuronal nitric oxide synthase (nNOS), were found to highly co-express TPH2 in the DRv (Fig. 3D,E) with 98.03 ± 1.44 % of nNOS+ neurons co-expressing TPH2 in non-stressed animals (Fig. 3E). There were no significant differences in the total number of nNOS+ neurons (Fig. 3I) or nNOS+ TPH2+ co-expressing neurons (Fig. 3J) across conditions. A 98 % TPH2+ nNOS+ co-expression indicated that there were not enough nNOS+ TPH2− neurons to account for the rise in TPH2+ numbers in susceptible rats. VGLUT3-expressing neurons have been shown to co-express TPH2, project to the VTA, and drive reward (Wang et al., 2019). Accordingly, we immunostained (Fig. 3F) and quantified the number of DRv glutamatergic neurons, marked by vesicular glutamate transporter isoform 3 (VGLUT3) and found that 86.51 ± 2.78 % of VGLUT3+ neurons co-express TPH2 in control brains (Fig. 3G). We hypothesized that the remaining VGLUT3+ TPH2− neurons could represent a significant portion of the reserve pool available for stress-induced TPH2 recruitment. To test whether VGLUT3 is a ‘switching partner’ for TPH2 as previously shown for TH (Dulcis and Spitzer, 2008; Dulcis et al., 2013), we quantified the number of total VGLUT3+ and VGLUT3+ TPH2+ co-expressing neurons in the DRv. A 1-way ANOVA (F(2,14)=7.001, *p*=0.008) revealed a significant decrease in the number of VGLUT3+ neurons in both susceptible (Tukey’s HSD: *p*=0.039) and resilient (Tukey’s HSD: *p*=0.007) rats relative to controls (Fig. 3H,K). The number of VGLUT3+ TPH2+ co-expressing neurons was also decreased in stressed rats (Fig. 3L, 1-way ANOVA F(2,14)=5.367, *p*=0.019) with a significant difference between resilient and control groups (Tukey’s HSD: *p*=0.023) and a strong trend toward a significant decrease in the susceptible group (Tukey’s HSD: *p*=0.054).

**Figure 3:**
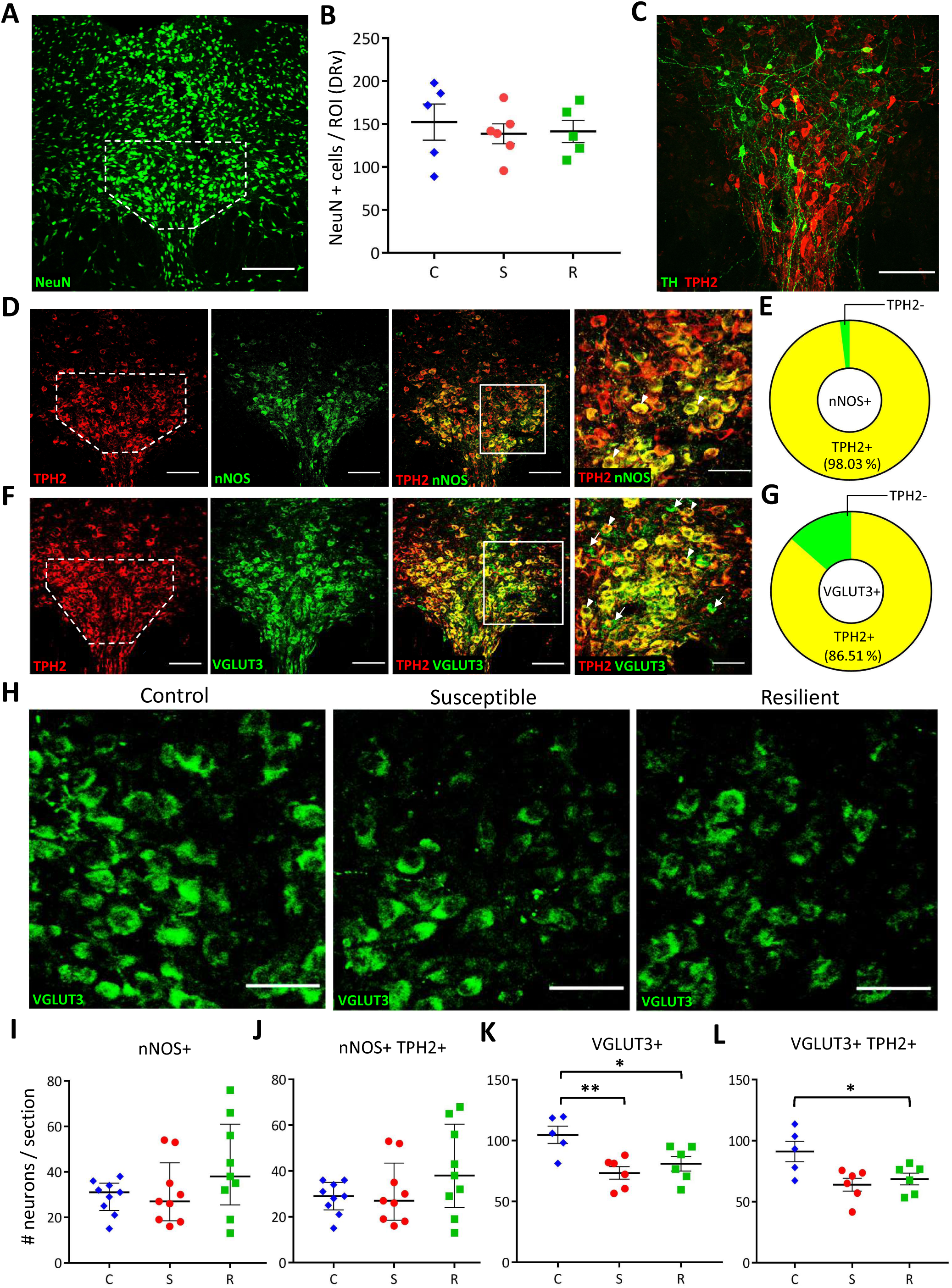
Stressed rats have fewer VGLUT3+ neurons in the DRv. **A**, Representative coronal DR section showing NeuN immunoreactivity. DRv margins were outlined in white. Scale bar: 200 µm. **B**, Quantification of NeuN+ cells in control (C, blue diamonds, *n=5 rats*), susceptible (S, red circles, *n=6 rats*), and resilient (R, green squares, *n=5 rats*) groups. ROI, region of interest. Counts were normalized to ROI area, averaged across rats per group and plotted as mean ± s.e.m. **C**, Representative coronal DR section showing dopaminergic (TH+, green) and serotonergic (TPH2+, red) but no co-expression. Scale bar: 100 µm. **D**, Representative images of a coronal DR section showing nitrergic (nNOS) neurons co-expressing TPH2. DRv outlined in white. Left to right: TPH2 (red), nNOS (green), merge (scale bar: 100 µm), and higher magnification of ROI drawn in merged image (scale bar: 25 µm). Arrowheads indicate nNOS+ TPH2+ (yellow) co-expressing neurons. **E**, Quantification of TPH2/nNOS co-expression in the DRv. Yellow sector indicates percentage of nNOS+ neurons co-expressing TPH2. Green sector indicates nNOS-only neurons. **F**, Representative images of coronal DR section showing glutamatergic (VGLUT3+) neurons co-expressing TPH2. DRv outlined in white. Left to right: TPH2 (red), VGLUT3+ (green), merge (scale bar: 100 µm), and higher magnification of ROI drawn in merged image (scale bar: 50 µm). Arrowheads indicate VGLUT3+ TPH2+ (yellow) co-expressing neurons. **G**, Quantification of TPH2/VGLUT3 co-expression in the DRv. Yellow sector indicates percentage of VGLUT3+ neurons also expressing TPH2. Green sector indicates VGLUT3-only neurons. **H**, VGLUT3+ neurons in the DRv of control, susceptible and resilient groups (scale bar 50 µm). **I**, **J**, Quantification of nNOS+ (**I**) and nNOS+ TPH2+ co-expressing **(J**) neurons in the DRv of control (C, blue diamonds), susceptible (S, red circles) and resilient (R, green squares). Counts were obtained from 3 sections each from 3 rats per group and plotted as median with interquartile range. **K**, Quantification of DRv VGLUT3+ neurons. **L**, Quantification of DRv VGLUT3+ TPH2+ co-expressing neurons. For **K** and **L**, counts were averaged across rats (Controls *n=5*, susceptible *n=6*, resilient *n=6*) and plotted as mean ± s.e.m. * *p* < 0.05, ** *p* < 0.01. Graph symbols as defined in **I**.

To test whether the observed changes in TPH2 and VGLUT3 protein arose from changes in the corresponding mRNA, we performed *in-situ* hybridization for *Tph2* and *Vglut3* mRNA (Fig. 4A). Additionally, we also probed for *Pet1* mRNA (Fig. 4A), which encodes a key transcription factor controlling serotonergic identity (Hendricks et al., 1999). PET1 regulates transcription of *Tph2*, *Sert* and other components of serotonergic identity (Spencer and Deneris, 2017). As expected, we found that all neurons expressing *Tph2* mRNA also expressed *Pet1* mRNA (Fig. 4C,D; arrowheads); however, Kruskal Wallis H tests revealed no significant differences across experimental groups in levels of *Tph2*, *Vglut3* or *Pet1* mRNA (Fig. 4B). Interestingly, a small fraction (9.17 ± 2.99 %) of *Pet1*-expressing neurons did not contain *Tph2* mRNA; a further subset (66.66 ± 11.33 %) of which co-expressed *Vglut3* mRNA (Fig. 4D; dashed outlines). The *Pet1+ Tph2−* neurons (both glutamatergic and non-glutamatergic) might represent an additional reserve pool, that is not recruited by 21-day social defeat, but might be primed for acquisition of *Tph2* mRNA and TPH2 protein when induced by prolonged chronic stress, extending beyond 3 weeks.

**Figure 4:**
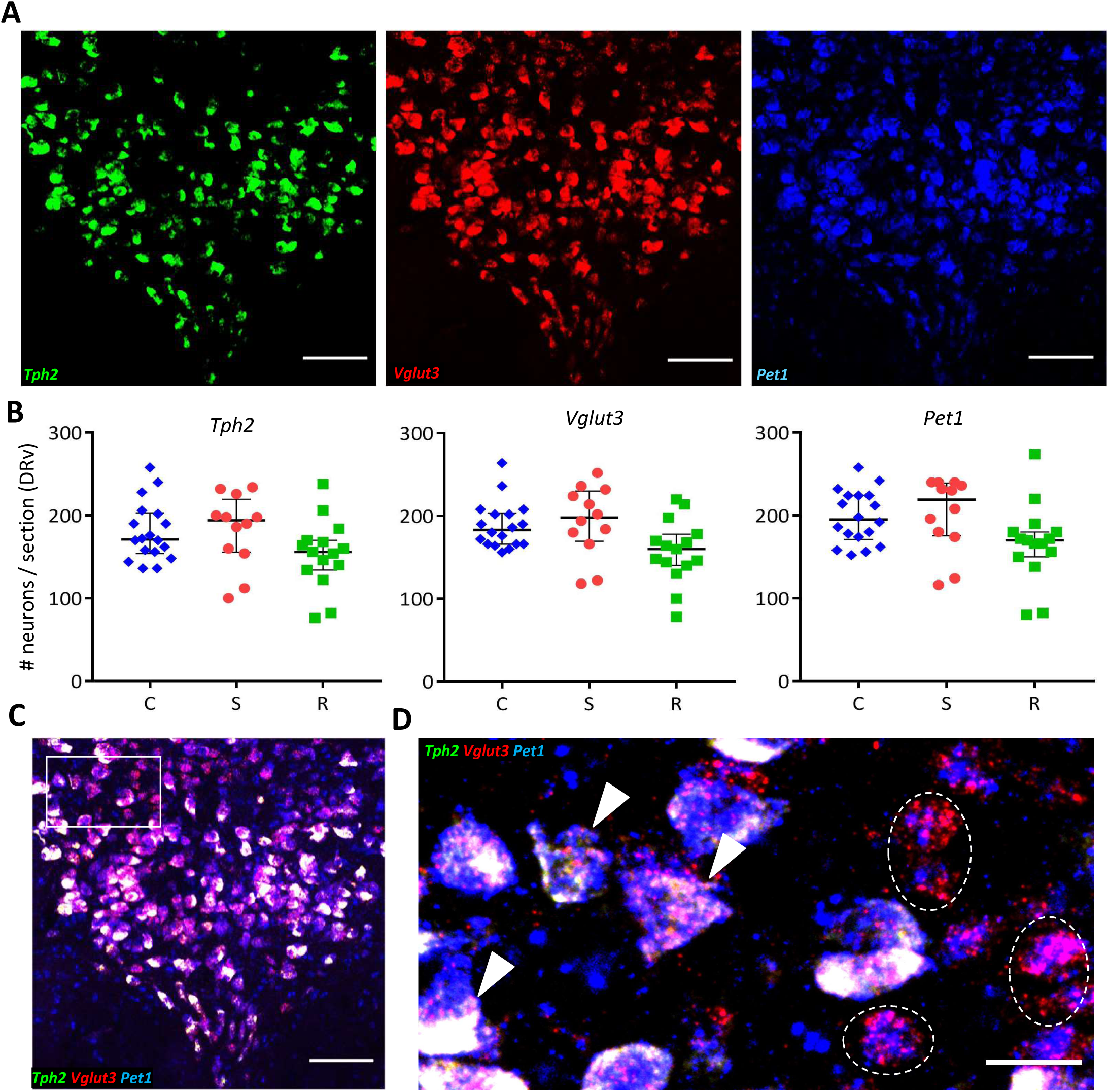
Levels of *Tph2*, *Vglut3* and *Pet1* mRNA do not differ across stress conditions. **A**, Representative images of a coronal DR section processed for *Tph2* (green), *Vglut3* (red) and *Pet1* (blue) triple *in* situ hybridization. Scale bars: 100 µm. **B**, Quantification of the number of neurons expressing *Tph2*, *Vglut3* and *Pet1* mRNA across control (C, blue diamonds, *n=5 rats*), susceptible (S, red circles, *n=4 rats*) and resilient (R, green squares, *n=4 rats*) groups. Counts were obtained from 1-3 DRv sections from each rat and plotted as median with interquartile range. **C**, Merge of 3 channels shown in **A**. Scale bar: 100 µm. **D**, ROI (rectangle in **C**) at higher magnification showing triple labelled (arrowheads), and non-serotonergic *Pet1+ Vglut3+ TPH2−* (dashed outline) neurons. Scale bar: 25µm.

### Stressed rats display lower cFos expression in DRv neurons

Neurotransmitter plasticity is activity-dependent (Borodinsky et al., 2004; Velazquez-Ulloa et al., 2011; Guemez-Gamboa et al., 2014; Meng et al., 2018). To investigate whether the recruitable reserve pool of TPH2− neurons in the DRv experiences any change in neuronal activity in response to social defeat, we assessed cFos expression by immunohistochemistry across stress conditions (Fig. 5A). The total number of cFos-immunoreactive neurons was decreased following stress (Fig. 5A,C; Kruskal Wallis H test, χ^2^(2)=11.325, *p*=0.003). *Post hoc* pairwise comparisons with Bonferroni correction showed that the decrease in cFos was significant in susceptible (*p*=0.004) and resilient (*p*=0.044) rats relative to controls. Specifically, this decrease occurred in TPH2− neurons (Fig. 5B,D; Kruskal Wallis H test, χ^2^(2)=17.967, *p*<0.001), both in susceptible (*p*<0.001) and resilient (*p*=0.031) rats, while cFos expression was unchanged in TPH2+ neurons (Fig. 5E). The observed differences in cFos expression across groups indicated that neuronal activity changed specifically in TPH2− DRv neurons in response to social stress. We next tested whether activity manipulation in these neurons results in alteration of their TPH2 phenotype, ultimately modulating behavior.

**Figure 5:**
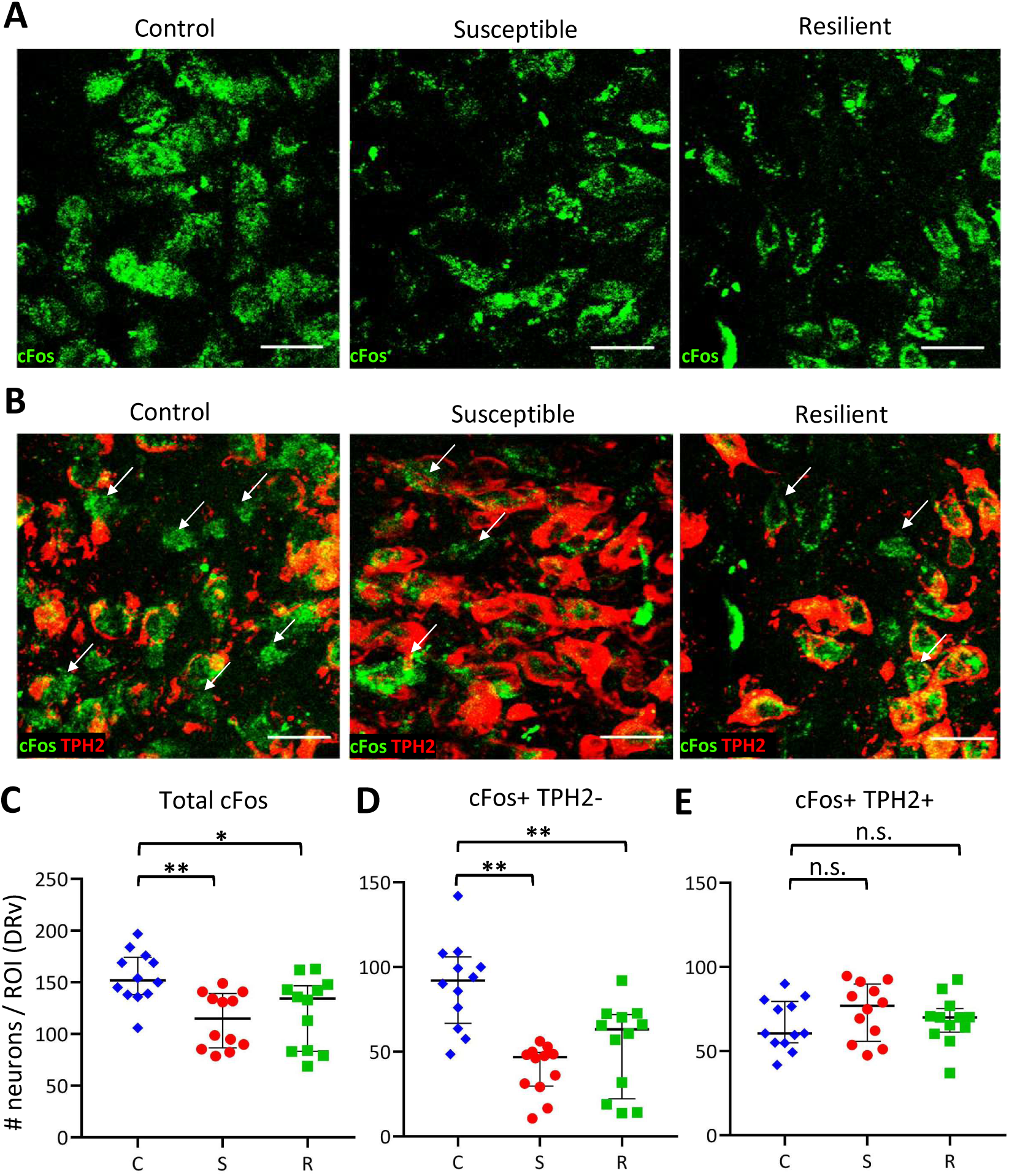
Stressed rats have fewer active TPH2− neurons in the DRv. **A**, cFos immunoreactivity in DRv of control, susceptible and resilient rats. Scale bars: 25 µm. **B**, cFos immunoreactivity in TPH2+ (red) and TPH2− (arrows) neurons in DRv across groups. Scale bars: 25 µm. **C**, **D**, **E**, Quantification of total cFos+ (**C**), cFos+ TPH2− (**D**), and cFos+ TPH2+ (**E**) neurons in controls (C, blue diamonds), susceptible (S, red circles), and resilient (R, green squares). Counts were obtained from 4 sections each from 3 rats per group, normalized to ROI area, and plotted as median and interquartile range. \**p* < 0.05, \*\**p* < 0.01, n.s. not significant (*p*>0.05).

### Chronic activation of amygdalar CRH+ neurons during stress ameliorates anhedonia and prevents TPH2 induction

To investigate whether TPH2 plasticity is activity-dependent and whether resilience is inducible in animals subjected to social stress, we manipulated neuronal activity in the DRv, by using DREADDs to drive the activity of one of its major inputs (Pollak Dorocic et al., 2014; Weissbourd et al., 2014), the central amygdala (CeA) (Fig. 6A). In *Crh-Cre* transgenic rats (Pomrenze et al., 2015), the CeA was bilaterally transfected with a Cre-dependent AAV vector encoding the excitatory DREADD receptor (hM3Dq) tagged with mCherry (Fig. 6A,B). A Cre-dependent AAV vector expressing GFP was used as a control (Fig. 6C). Rats were then trained and baselined in the ICSS procedure, habituated to intraperitoneal injections of saline and tested for anhedonia during 21-day social defeat with daily clozapine pretreatment. One-way repeated measures ANOVA with Greenhouse-Geiser correction revealed that saline injections did not significantly alter reward thresholds (Fig. 6D). To test whether clozapine by itself affected reward thresholds relative to baseline over the 21-day period, the non-stressed control group expressing GFP virus was examined using a 1-way repeated measures ANOVA with Greenhouse-Geiser correction. The within-subjects effect of *Day* was not significant (Fig. 6E). To test whether DREADD-mediated activation of the CeA affected reward thresholds in the absence of stress, thresholds of non-stressed controls expressing DREADDs and treated with clozapine for the 21-day period, were analyzed similarly. No significant effect of *Day* was observed (Fig. 6E), indicating that any effect of DREADD activation on reward thresholds in stressed rats was specific to the stress response and not a general reward-enhancing or reward-diminishing effect of stimulating CRH neurons of the CeA. A 2-way mixed ANOVA with Greenhouse-Geiser correction revealed that the two groups of non-stressed controls (GFP or hM3Dq virus injected) did not differ significantly from each other. Therefore, the two control groups were combined and used as a pooled control group for all subsequent analyses. Based on previous evidence that susceptibility or resilience manifests after *chronic,* but not *acute*, stress (Der-Avakian et al., 2014) which was supported by our observations (Fig. 1A,B), we analyzed the acute (days 1-3) and chronic (days 19-21) effects of social defeat on reward thresholds using a 2-way mixed ANOVA (no violation of sphericity assumption, ε=1), with *Period* (acute or chronic) as the within-subjects factor and *Group* as the between-subjects factor (Fig. 6F). There was a significant interaction of *Period* × *Group* (F(4,24)=2.994, *p* = 0.039) and significant main effects of *Period* (F(1,24)=5.329, *p* = 0.030) and *Group* (F(4,24)=16.922, *p* = 1.0E-6) (Fig. 6F).

**Figure 6:**
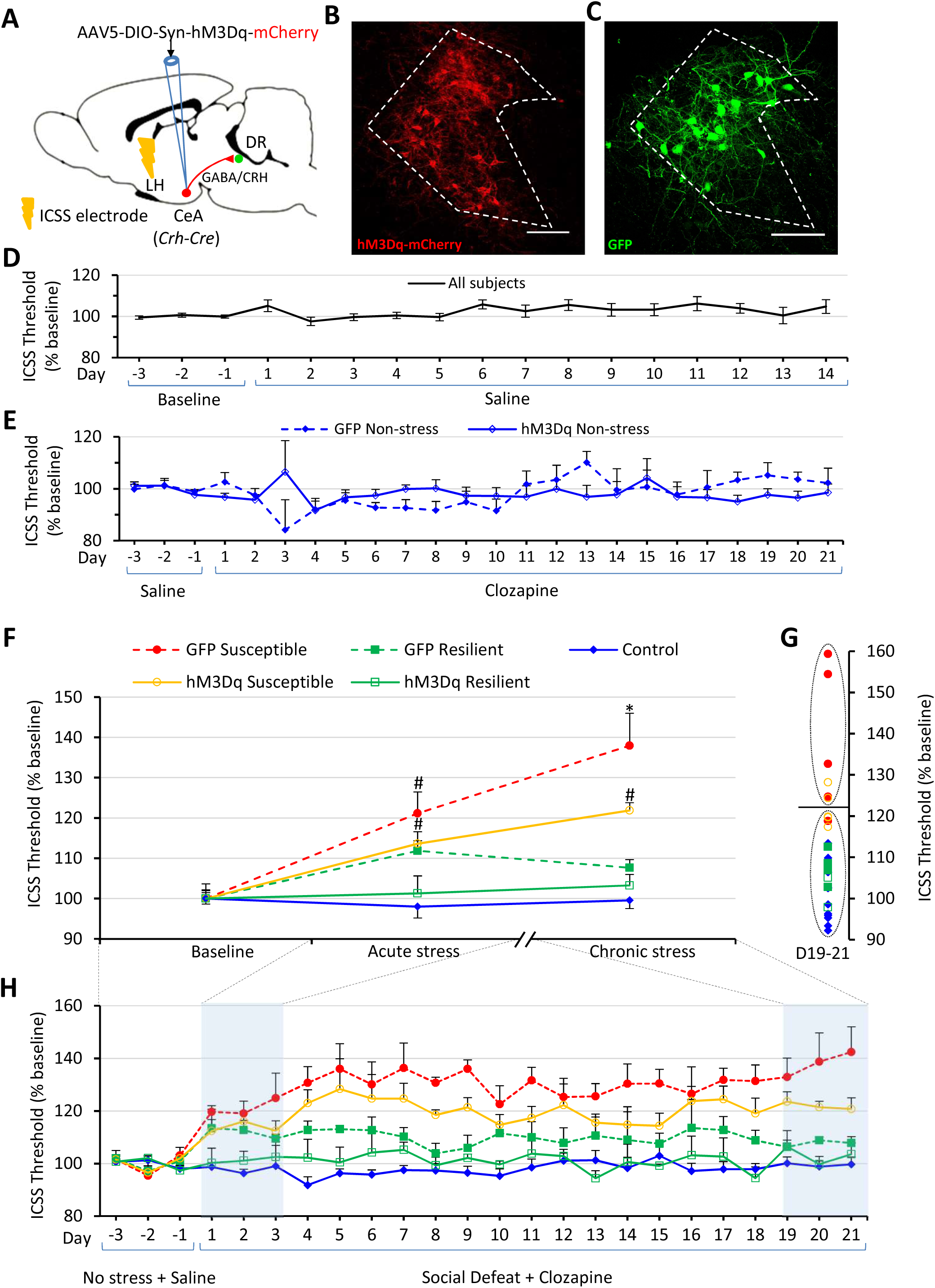
Chronic activation of amygdalar CRH+ neurons reduces stress-induced reward threshold elevations. **A**, Cartoon showing viral strategy to chemogenetically activate CRH/GABA neurons of the central amygdala (CeA) in *Crh-Cre* rats. LH, lateral hypothalamus; DR, dorsal raphe nucleus. **B**, Expression of Cre-dependent mCherry-tagged excitatory DREADD (hM3Dq) virus in a *Crh-Cre* rat. Cell bodies and fibers localized to the lateral subnucleus of the central amygdala (CeL, dashed outline) are visible in red. **C**, Expression of Cre-dependent GFP virus in CeL (dashed outline) of a *Crh-Cre* rat. Scale bars in **B**,**C**: 100 µm. **D**, Daily ICSS thresholds (mean across rats ± s.e.m.) plotted as percent of baseline for 3 days prior to saline IP injections and each day for saline treatment. All subjects (*n=29 rats*) are plotted. **E**, Daily ICSS thresholds (mean across rats + s.e.m.) plotted as percent of baseline, for 3 days of baseline prior to clozapine treatment and each day of 21-day clozapine treatment in non-stressed controls. GFP-expressing rats (solid diamonds, dotted line, *n=4*) and hM3Dq-expressing rats (hollow diamonds, solid line, *n=*8) are plotted. **F**, ICSS thresholds plotted as percent of baseline (mean across rats +/- s.e.m.) for baseline (average of days -3 to -1) with saline treatment, acute (average of days 1-3) and chronic (average of days 19-21) social defeat with clozapine treatment. GFP-expressing susceptible rats (red solid circles, dotted line, *n=5*), GFP-expressing resilient rats (green solid squares, dotted line, *n=4*), hM3Dq-expressing susceptible rats (orange hollow circles, solid line, *n=5*) and hM3Dq-expressing resilient rats (green hollow squares, solid line, *n=3*) are plotted. Significant (*p*<0.05) post-hoc pairwise comparisons for each timepoint are indicated above error bars. # indicates significant difference relative to control but not to other groups. * indicates significant difference relative to control and resilient groups expressing the same virus. **G**, k-means cluster analysis of Days 19-21 averaged ICSS thresholds. Sample sizes and graph symbols as in **F**. **H**, Effect of DREADD stimulation of CeA CRH+ neurons on ICSS thresholds during social defeat. Daily ICSS thresholds (mean across rats + s.e.m.) plotted as percent of baseline, for 3 days of baseline prior to clozapine treatment/social defeat and each day of 21-day clozapine treatment/social defeat. Graph symbols as in **F**. Days used to compute averages for acute and chronic stress in **F** are highlighted in blue.

Pairwise *post hoc* comparisons with Bonferroni correction showed that GFP susceptible and hM3Dq susceptible rats had significantly elevated reward thresholds relative to non-stressed controls after acute stress exposure (GFP susceptible: *p*=0.001, hM3Dq susceptible: *p*=0.032). However, as hypothesized, the effects of chronic stress differed from those of acute stress across groups. After chronic stress (days 19-21), while the reward thresholds of GFP susceptible rats were significantly elevated relative to controls (*p*=5.51E-7) and GFP resilient rats (*p=*0.001), the reward thresholds of hM3Dq susceptible rats were not significantly elevated relative to the hM3Dq resilient group (*p*=0.115), but only elevated relative to control (*p*=0.001). In other words, the reward thresholds of the DREADD-treated susceptible rats were statistically similar to the resilient group, suggesting that DREADD-activation of CeA CRH+ neurons ameliorated chronic stress-induced anhedonia. A k-means cluster analysis (based on averaged thresholds of days 19-21) classified all controls and resilient rats into a single cluster (Fig. 6G). This cluster also included 3 out of 5 hM3Dq susceptible, and 1 out of 5 GFP susceptible rats. Other rats from the GFP susceptible and hM3Dq susceptible groups formed a separate cluster. Extended (day-wise) reward thresholds during chemogenetic manipulation are shown in Figure 6H.

At the anatomical level, the number of DRv TPH2+ neurons showed significant differences across groups (Fig. 7A; F(4,24)=6.337, *p*=0.001). *Post hoc* pairwise comparisons showed that the GFP susceptible group had a significantly elevated number of TPH2+ neurons relative to controls (Tukey’s HSD: *p*=0.041), hM3Dq resilient (Tukey’s HSD: *p*=0.014) and GFP resilient (Tukey’s HSD: *p*=0.001) groups, recapitulating the effects of social defeat in wildtype rats (Fig. 2D) and demonstrating that clozapine alone had no effect on neurotransmitter plasticity in the DR following social defeat. Interestingly, the hM3Dq susceptible group did not display an elevated number of TPH2+ neurons relative to control (Tukey’s HSD: *p*=9.99E-1), hM3Dq resilient (Tukey’s HSD: *p*=0.620), or GFP resilient (Tukey’s HSD: *p*=0.142) groups, indicating that manipulation of DR activity via CeA activation prevented stress-induced increase in TPH2+ neuron number. The lower extent of anhedonia in the hM3Dq susceptible rats, compared to GFP susceptible rats (Fig. 6F,G) is reflected in the lower TPH2+ neuron numbers in the hM3Dq susceptible group (Fig. 7A). TPH2+ neuron number in the DRv and ICSS thresholds were significantly positively correlated in all stressed rats (wildtype and transgenic) used in this study (Fig. 7B; Pearson’s r=0.743, *p*=3.0E-6, *n*=30). This suggests that the number of TPH2+ neurons in the DRv is a molecular marker of susceptibility to anhedonia induced by chronic stress in rats.

**Figure 7:**
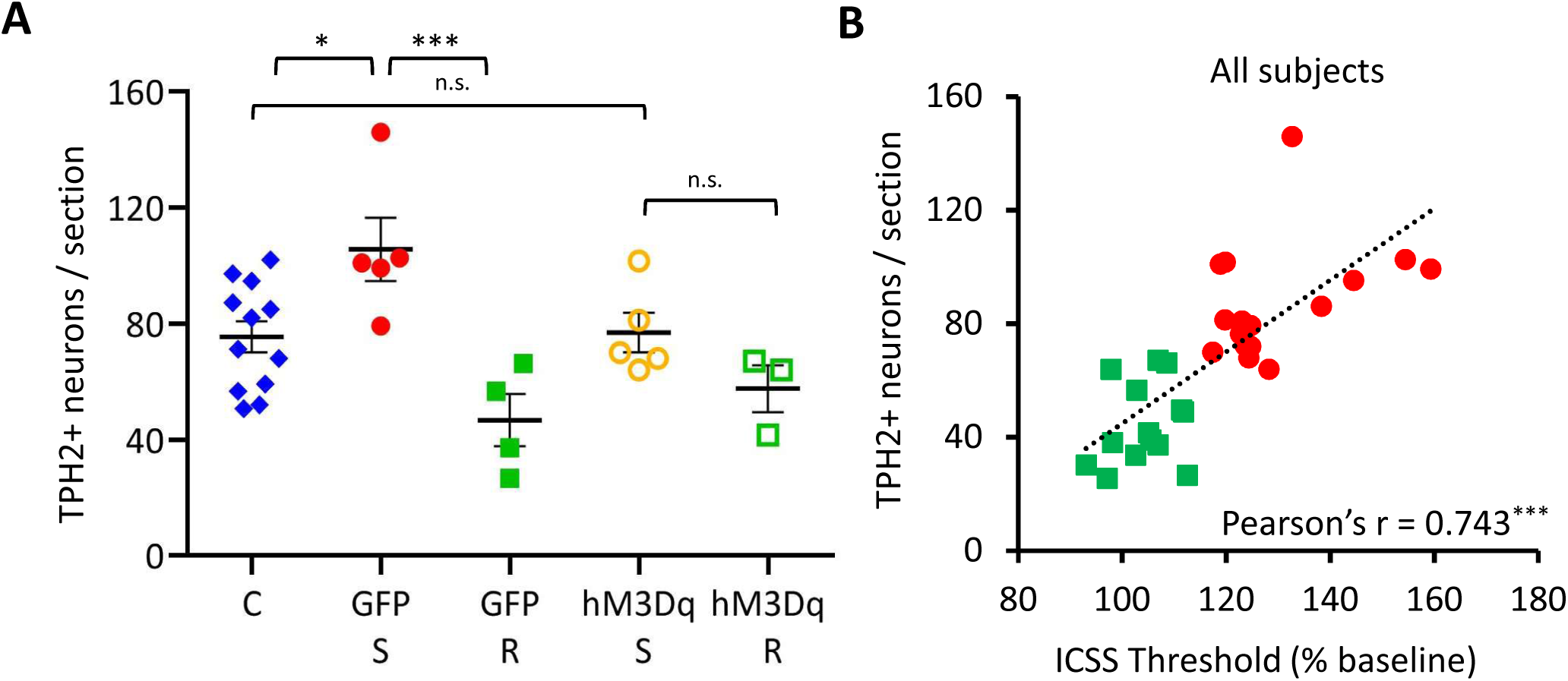
Chronic activation of amygdalar CRH+ neurons abolishes stress-induced gain of TPH2 in susceptible rats. **A**, Quantification of TPH2+ neurons in the DRv from pooled control (C, blue diamonds, *n=12 rats*), GFP-expressing susceptible (GFP-S, red solid circles, *n=5 rats*), GFP-expressing resilient (GFP-R, green solid squares, *n=4 rats*), hM3Dq-expressing susceptible (hM3Dq-S, orange hollow circles, *n=5 rats*) and hM3Dq-expressing resilient (hM3Dq-R, green hollow squares, *n=3 rats*) groups. Counts were averaged across rats and plotted as mean ± s.e.m. * *p* < 0.05, *** *p* < 0.001, n.s. not significant (*p* > 0.05). **B**, Correlation analysis of DRv TPH2+ neurons and ICSS thresholds (Average of Days 19-21) for all stressed rats (wildtype and *Crh-Cre*) used in this study. Rats were classified as susceptible (red solid circles, *n=16*) or resilient (green solid squares, *n=14*), across genotypes and virus groups. Pearson’s product-moment correlation coefficient shown on bottom right for each analysis. *** *p* < 0.001.

## DISCUSSION

Previous studies in rodent models have shown that serotonergic molecular machinery is upregulated in the DR after both chronic (Adell et al., 1988; Zhang et al., 2012; Donner et al., 2016) and acute stress (Donner et al., 2018). However, these studies did not specifically address differences between animals that are susceptible or resilient to chronic stress-induced anhedonia, or whether this upregulation occurs in identified classes of neurons expressing specific neurotransmitters. Other molecular adaptations investigated in the context of susceptibility/resilience such as synaptic excitability in the nucleus accumbens (Krishnan et al., 2007; Vialou et al., 2010), VTA (Krishnan et al., 2007), medial prefrontal cortex (Covington et al., 2010; Lehmann and Herkenham, 2011) and HPA axis activation (Elliott et al., 2010) have not been demonstrated to be specific to anhedonia. Furthermore, neurotransmitter plasticity (Dulcis and Spitzer, 2012) as a mechanism to explain susceptibility/resilience to anhedonia has not been previously explored.

Our results revealed that chronic social defeat induced an increase in the number of TPH2+ neurons without affecting the total number of DRv neurons in susceptible rats, indicating that differentiated TPH2− neurons in the DR are recruited to additionally express TPH2. A concomitant reduction in VGLUT3+ neurons in the DRv of all stressed rats suggests that stress-induced anhedonia in susceptible animals is an example of transmitter switching driven by stress. Our finding is the first instance of a molecular and cellular marker based on neurotransmitter phenotype to be associated with vulnerability to stress. Neurotransmitter plasticity, also referred to as neurotransmitter switching (Dulcis & Spitzer, 2012), has been shown to occur in response to other forms of stress such as altered photoperiod exposure (Dulcis et al., 2013; Young et al., 2018) or exposure to psychostimulants such as nicotine (Romoli et al., 2019) and methamphetamine (Kesby et al., 2017). Interestingly, the change in number of TPH2+ neurons occurred exclusively in the DRv, but not in the other subnuclei of the mid-rostrocaudal DR. This is consistent with what is known about the afferent and efferent connectivity of the DRv. The DRv receives inputs from the lateral hypothalamus, central amygdala and orbito-frontal cortex (Peyron et al., 1997) and projects to orbito-frontal and other cortical areas (Ren et al., 2018) and the VTA (Qi et al., 2014). All of these areas are implicated either in reward sensing and integration (Rolls, 2000; Gottfried et al., 2003; van Zessen et al., 2012), reward-based decision making (Bechara et al., 2000), or stress processing (Swiergiel et al., 1993; Hsu et al., 1998; Bonnavion et al., 2015). The DRv is therefore, a critical hub that regulates stress and reward-related behaviors by serotonergic neuromodulation of multiple target areas. Neurotransmitter plasticity in the DRv could impact the activity and function of these regions, subsequently affecting behavior.

The increase in number of TPH2+ neurons in the anhedonic condition (susceptible animals), was somewhat unexpected, given that serotonergic depletion is conventionally associated with depression based on studies that used SSRI treatment (Delgado, 2004; Abumaria et al., 2007), tryptophan treatment (Persson, 1967; Glassman and Platman, 1969; Wålinder et al., 2003) or 5-HT depletion (Shopsin et al., 1976; Delgado et al., 1994; Sachs et al., 2015). However, increased TPH2 immunoreactivity has been observed after exposure to stress (Poeggel et al., 2003) and in brains of depressed patients who committed suicide (Underwood et al., 1999; Boldrini et al., 2005), suggesting that this may be a homeostatic mechanism by the DR to compensate for serotonin depletion and deficient signaling in target areas. There is also ample electrophysiological evidence to suggest that serotonin exerts complex neuromodulation of the VTA in conjunction with glutamate and dopamine (Pessia et al., 1994; Gervais and Rouillard, 2000; Guiard et al., 2008; Liu et al., 2014; Belmer et al., 2016; Wang et al., 2019); therefore an increase in TPH2+ neurons in the DRv may lead to reduced reward function in the VTA.

The opposing regulation of TPH2 and VGLUT3 observed in susceptible animals and the parallel finding that some VGLUT3+ neurons already express *Pet1* transcripts in unstressed animals, suggest that this glutamatergic pool might be primed for TPH2 recruitment, similar to the mechanism described for nicotine-induced dopaminergic plasticity within a Nurr1-expressing reserve pool in the VTA (Romoli et al., 2019). Since the number of neurons gaining TPH2 in susceptible animals is greater than the number of neurons losing VGLUT3, the existence of additional non-serotonergic neurons comprising the total reserve pool is expected. A fraction of GABA-expressing neurons in DRv (Fu et al., 2010; Bang and Commons, 2012) could be one such additional reserve pool. The absence of changes in *Tph2 or Vglut3* at the transcript level (Fig. 4) suggests that the observed differences in the number of TPH2+ and VGLUT3+ neurons across groups (Fig. 3) arose from differences at the level of post-transcriptional or translational regulation similar to miRNA-mediated dopamine/GABA switching in response to social cues in the developing amphibian olfactory bulb (Dulcis et al., 2017).

Neuronal activity, as measured by cFos expression, was decreased specifically in DRv TPH2− neurons following chronic stress, suggesting DRv inhibition after chronic stress. This is consistent with previous observations of serotonergic induction following inhibition of neuronal activity (Demarque and Spitzer, 2010). The DRv receives projections from the central amygdala (CeA) containing GABA and corticotropin releasing hormone (CRH), which modulate the activity of both serotonergic and non-serotonergic neurons (Pollak Dorocic et al., 2014; Weissbourd et al., 2014; Pomrenze et al., 2015) in the DR, via CRHR1 and CRHR2 receptors (Lowry et al., 2000; Day et al., 2004; Donner et al., 2016). Moreover, the CeA responds differently to acute versus chronic social defeat (Martinez et al., 1998, 2002a). Further, CRH in particular is implicated in the development of depression-like phenotypes in rodents in response to uncontrollable stressors (Maier and Watkins, 2005). These reports which clearly indicated that the CRH+ neurons in the CeA form a strong input to the DRv involved in regulation of stress and reward-related behavior, motivated our choice to manipulate their activity. Lack of a significant difference between hM3Dq-treated susceptible and resilient rats at the end of chronic social defeat (days 19-21), suggested an antidepressant effect of activating CeA CRH+ neurons. It is likely that other brain regions involved in stress-processing and resilience, such as the medial prefrontal cortex (Amat et al., 2005), play a role in inducing resilience. Simultaneous activation of such regions might result in stronger effects on reward thresholds and resilience than what was achieved with CeA manipulation alone. The lack of a significant increase in the number of DRv TPH2+ neurons observed in hM3Dq-treated rats exposed to social defeat revealed the activity-dependent nature of resilience and its association with neurotransmitter phenotype in the DRv.

In conclusion, our findings begin to reveal a possible model of neurotransmitter plasticity involved in the regulation of resilience to social stress (Fig. 8). Susceptible animals gain TPH2 expression in the DRv in response to chronic stress, while resilient animals do not. VGLUT3 is lost in susceptible and resilient animals, largely by neurons that co-express both markers. This plasticity of expression occurs at the protein level, presumably by post-transcriptional or translational regulation since the number of neurons making the corresponding mRNA transcripts was unchanged. Activation of CRH+ neurons in the CeA, which form a major input to the DR, modulates the effects of social stress on DR activity by preventing TPH2+ induction. This consequently prevents or ameliorates anhedonia, promoting resilience.

**Figure 8:**
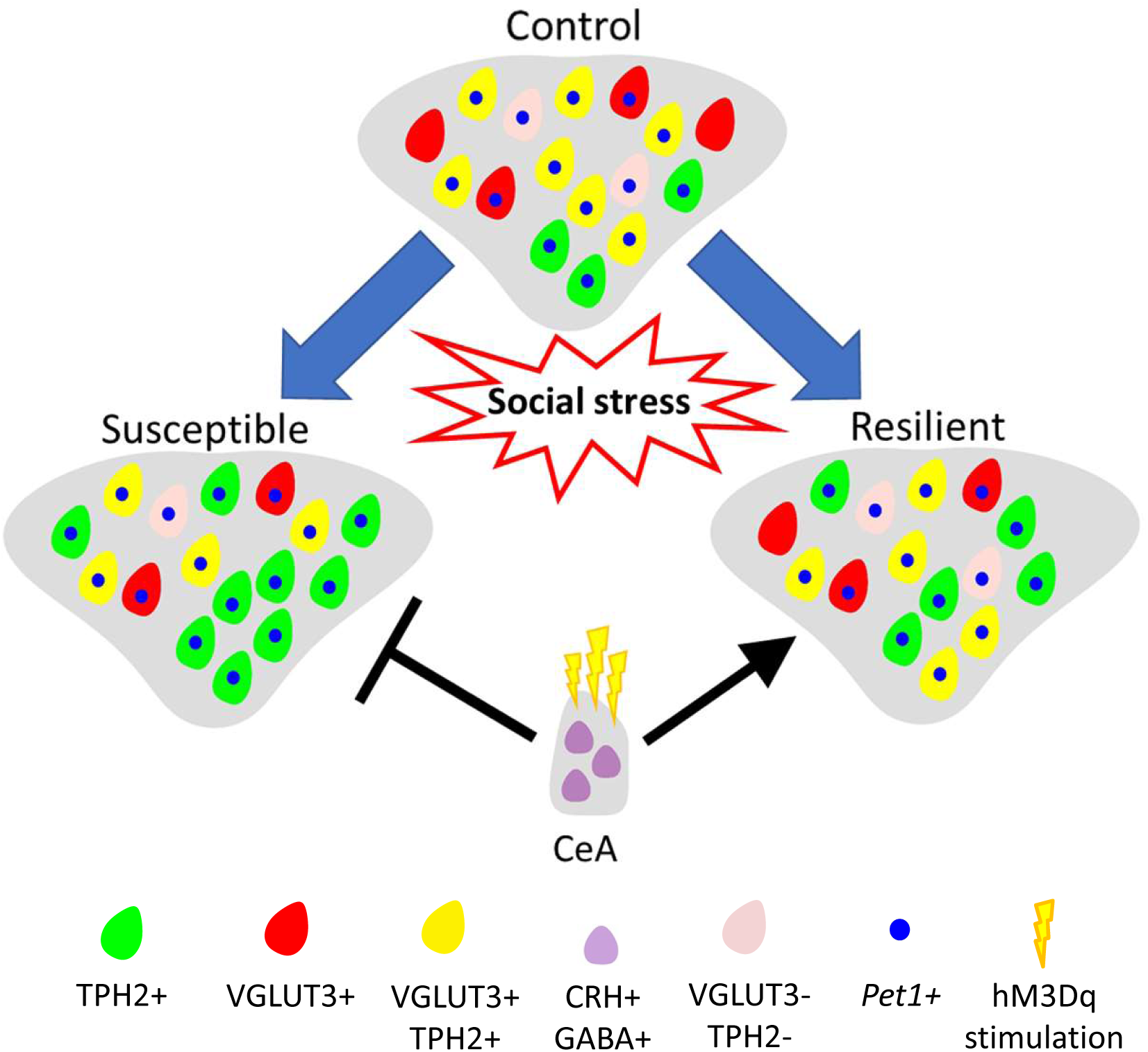
Model of neurotransmitter plasticity in the DRv in response to chronic social stress. In response to chronic social stress, neurotransmitter plasticity occurs in the DRv of stressed rats. Rats susceptible to stress-induced anhedonia gain TPH2 and lose VGLUT3 while resilient rats only lose VGLUT3. Loss of VGLUT3 leads to lower co-expression of the two markers in both conditions. The plasticity occurs in neurons expressing *Pet1* transcripts. Activation of CRH+ neurons of the CeA promotes resilience and blocks susceptibility.

Knowledge of the molecular signature of the reserve pool neurons that are recruited to a TPH2 phenotype in animals susceptible to chronic-stress and the activity-dependent induction of resilience could both be harnessed in the future to develop novel treatment strategies to elicit resilience and ameliorate stress-related disorders.

## Acknowledgements

This work was supported by the following grants: the Kavli Institute for Brain and Mind (Grant No. 2012-008 [to DD]), the Tobacco-Related Disease Research Program (Grant No. 27IR-0020 [to DD]), the National Institute of Drug Addiction (Grant No. R21-DA047455 [to DD]), the National Institute of Mental Health (Grant No. R01 MH106865 [to AD]), and the National Institute on Alcohol Abuse and Alcoholism (Grant No. R01 AA026560 [to AD]).

We thank Dr. Robert O. Messing, University of Texas, Austin, for kindly gifting us breeding pairs for the Crh-Cre transgenic rat line. We would also like to thank Dr. Christina M. Gremel, Dr. Thomas S. Hnasko and Dr. Samuel Barnes for their critical feedback on the research and Dr. Byung Kook Lim for his valuable feedback and the use of his slide-scanning microscope.

